# Integrated analysis of single-cell and transcriptome based RNA-seq multilayer network and WGCNA for construction and validation of an immune cell-related prognostic model in clear cell renal cell carcinoma

**DOI:** 10.1101/2021.10.15.464475

**Authors:** Guanlin Wu, Weiming Guo, Shuai Zhu, Gang Fan

## Abstract

Clear cell renal cell carcinoma (ccRCC) is the most common type of renal cancer (RCC). The increasing incidence and poor prognosis of ccRCC after tumour metastasis makes the study of its pathogenesis extremely important. Traditional studies mostly focus on the regulation of ccRCC by single gene, while ignoring the impact of tumour heterogeneity on disease progression. The purpose of this study is to construct a prognostic risk model for ccRCC by analysing the differential marker genes related to immune cells in the single-cell database for providing help in clinical diagnosis and targeted therapy. Single-cell data and ligand-receptor relationship pair data were downloaded from related publications, and ccRCC phenotype and expression profile data were downloaded from TCGA and CPTAC. The DEGs and marker genes of the immune cell were combined and then intersected with the ligand-receptor gene data, and the 981 ligand-receptor relationship pairs obtained were intersected with the target gene of the transcription factor afterwards; 7,987 immune cell multilayer network relationship pairs were finally observed. Then, the genes in the network and the genes in TCGA were intersected to obtain 966 genes for constructing a co-expression network. Subsequently, 53 genes in black and magenta modules related to prognosis were screened by WGCNA for subsequent analysis. Whereafter, using the data of TCGA, 28 genes with significant prognostic differences were screened out through univariate Cox regression analysis. After that, LASSO regression analysis of these genes was performed to obtain a prognostic risk scoring model containing 16 genes, and CPTAC data showed that the effectiveness of this model was good. The results of correlation analysis between the risk score and other clinical factors showed that age, grade, M, T, stage and risk score were all significantly different (p < 0.05), and the results of prognostic accuracy also reached the threshold of qualification. Combined with clinical information, univariate and multivariate Cox regression analyses verified that risk score was an independent prognostic factor (p < 0.05). A nomogram constructed based on a predictive model for predicting the overall survival was established, and internal validation performed well. Our findings suggest that the predictive model built based on the immune cells scRNA-seq will enable us to judge the prognosis of patients with ccRCC and provide more accurate directions for basic relevant research and clinical practice.

## Main content

ccRCC single-cell transcriptome data was downloaded from [1]. After standardized data processing, we obtained a total of 23 subgroup clusters, among which immune cells were mainly concentrated in subgroups 0, 1, 2, 3, 4, 5, 6, 7, 9, 12, 13, 15, 16, 18 and 22 **(Fig. S1, Table S1)**. Differentially expressed genes (DEGs) in the 23 clusters were listed in **(Fig. S2A, Table S2)**. The functional status of differentially expressed genes (DEGs) in immune cells were also performed **(Fig. S2B-F)**. Furthermore, a total of 42 immune cell marker genes related to ccRCC were downloaded from the CellMarker (http://bio-bigdata.hrbmu.edu.cn/CellMarker/) database and [2] performed differential expression analysis **(Fig. S2G)**.

In order to construct the ligand-receptor network, we first took the union of the differential genes of all immune cell clusters and the marker genes of all these immune cells. Afterwards, we intersected them with the ligand-receptor relationship pairs downloaded from [3]. Finally, a total of 981 pairs of ligand-receptor relationships were obtained **(Fig. S3A, Table S3)**. Afterwards, we downloaded the transcription factors and target genes data from the TRRUST (https://www.grnpedia.org/trrust/) database and merged the ligand-receptor network data with the transcription factors and target genes data, which is the intersection of the target genes of the transcription factors and the genes from the ligand-receptor network, we obtained 7,987 immune cell multifactor network relationship pairs **(Table S4)**. Then, 966 genes used for the immune cell multifactor network construction **(Fig. S3B)** were obtained by intersecting the 973 genes contained in the ligand-receptor network and the genes in TCGA (https://portal.gdc.cancer.gov/) dataset about ccRCC **(Table S5)**.

Through the WGCNA algorithm [4], the 966 genes in the immune cell multifactor network were used to construct a co-expression network to find the key modules related to overall survival (OS) and OS time. The appropriate scale-free power value was screened out as 4 **(Fig. S4A)** when the average connectivity degree was relatively higher [5]. All constructed modules are painted with different colours, and the cluster trees of genes are also shown in **Fig. S4B**. The black and magenta modules were chosen as the key modules, since they had the highest correlations with OS and OS time of ccRCC **(Fig. S4C-D)**. The correlations between module eigengenes (MEs) and clinic traits are shown in **Fig. S4E**. There were 53 genes in these two modules **(Table S6)**. For a deeper understanding about the biofunctions of these modules, genes in these modules were carried out to conduct gene ontology (GO) and Kyoto Encyclopedia of Genes and Genomes (KEGG) pathway analyses, the top terms were visualized **(Fig. S4F-I)**.

Using the “survival” R package to perform univariate Cox regression analysis on the 53 genes contained in the key modules of WGCNA, 28 genes with *P* value < 0.05 were obtained. Then, the 28 genes with significant prognostic differences were subjected to LASSO regression analysis. The specific steps were to first adopt the Cox proportional hazard model (family = “Cox”) to calculate the hazard ratio (HR) and *P* values of these genes **(Fig. 1A)** and then randomly simulated 1,000 times (maxit = 1000) to find the most suitable λ value **(Fig. 1B)**. Finally, “lambda.min” was used to screen out 16 genes for constructing a risk scoring model from these 28 genes **(Fig. 1C)**.

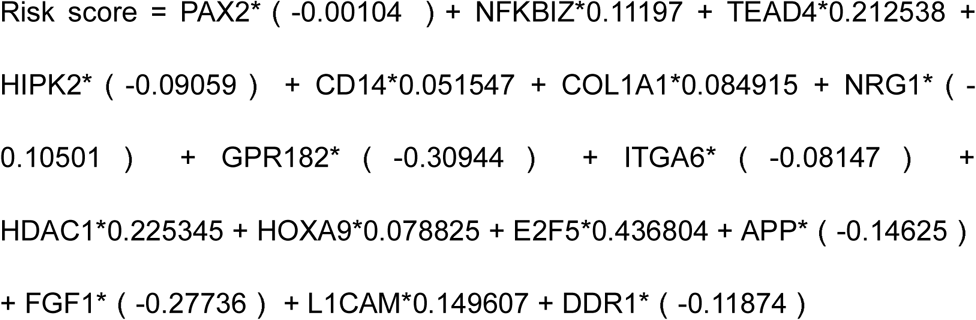

**Figure 1.**
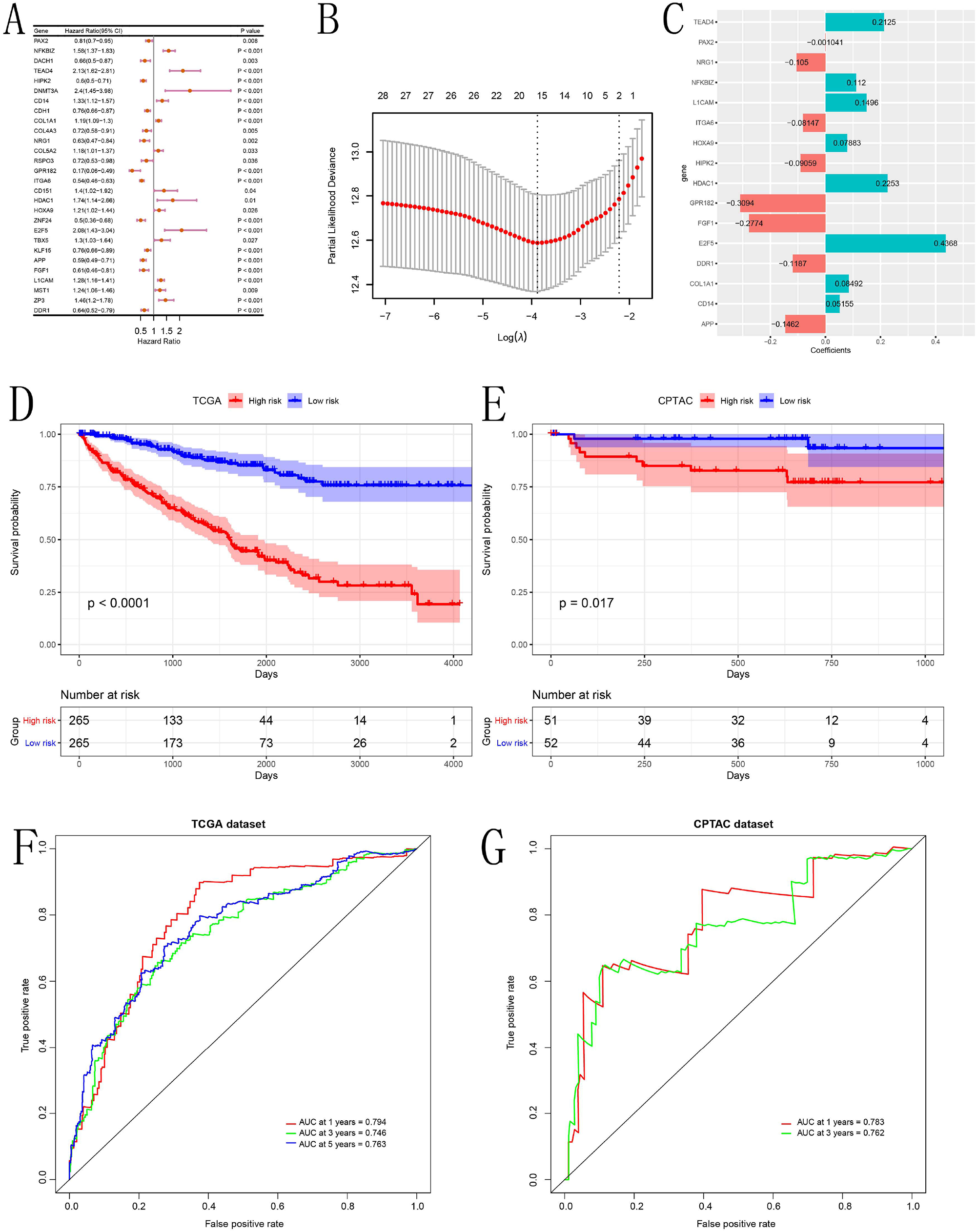
Construction and assessment of disease prognosis risk model for ccRCC. **(A)** Forest plot for univariate regression analysis of 28 genes. **(B)** Select the appropriate λ value through LASSO regression analysis. **(C)** Scoring chart of risk model constructed by 16 genes. **(D, F)** Evaluation of predictive effectiveness of risk prognostic model. **(E, G)** Use of the data in the CPTAC database as an external dataset for verification.

According to the risk score of each patient given by the model, the median was taken as the cut-off value, and the samples were divided into high and low risk groups. The time-dependent receiver operating characteristic (ROC) curve was used to evaluate the predictive ability of the model’s 1-, 3- and 5-year survival periods, and the survival curves of the high and low risk groups were also analysed. The CPTAC (https://cptac-data-portal.georgetown.edu/study-summary/S050) dataset for ccRCC was taken as the external validation database to verify the prognostic risk model. In the evaluation of the predictive efficacy of the prognosis model, Kaplan-Meier (K-M) survival analysis was performed on the high and low risk groups, and the difference was significant **(Fig. 1D)**. In the external CPTAC dataset, K-M survival analysis was performed on the high and low risk groups, and the difference was also significant **(Fig. 1E)**. Moreover, the AUC values in their ROC curves validated their precision **(Fig. 1F-G)**.

In the correlation comparison between risk score and other clinical factors, the results showed that all factors other than age had a certain correlation with risk score **(Fig. S5A-G)**. From the ROC curve, it was also found that, except for the N stage, the AUC values of other ROC curves were above 0.5, and the AUC values of the risk score and stage were above 0.7 **(Fig. S5H-J)**.

The immune infiltration of genes in the model is shown in **(Fig. 6A)**. The differential expression of them in TCGA **(Fig. 6B)** and CPTAC **(Fig. 6C)** are displayed. Univariate Cox regression analysis **(Fig. S6D)** and multivariate Cox regression analysis **(Fig. S6E)** for the independent analysis of the prognostic model were also done.

Additionally, we also developed a nomogram with predictors included age, gender, T, N, M, grade, risk score and stage **(Fig. S7A)**. The consistency index (C-index), which represents the discrimination, of the nomogram was 0.799. and our calibration chart revealed that the nomogram has a good predictive ability **(Fig. S7B-D)**. The AUC values indicated that the predictive precision of the nomogram combined with all factors was better **(Fig. S7E-G)**.

In summary, our research reveals that the immune cells in ccRCC have a significant impact on the prognosis of patients, and have obvious potential regulatory effects on the occurrence and development of ccRCC. The predictive model based on immune cell scRNA-seq will enable us to judge the prognosis of ccRCC patients and provide valuable guidance for clinical practice.

## Abbreviation

RCC: renal cell carcinoma
ccRCC: clear cell renal cell carcinoma
OS: overall survival
scRNA-seq: single-cell: RNA-sequencing
WGCNA: weighted gene co-expression network analysis
TCGA: The Cancer Genome Atlas
CPTAC: Consortium for Clinical Proteome Cancer Analysis
DEGs: differentially expressed genes
MEs: module eigengenes
GO: gene ontology
KEGG: Kyoto Encyclopedia of Genes and Genomes
HR: hazard ratio
ROC: receiver operating characteristic
C-index: concordance index
K-M: KaplanMeier

## Declarations

### Ethics approval and consent to participate

Not applicable.

### Consent for publication

Not applicable.

### Availability of data and material

The data and materials can be obtained by contacting the corresponding author.

### Authors’ contributions

G.W designed the research plan, analyzed datasets and wrote the manuscript. W.G and S.Z provided meaningful discussion of key points. G.F gave guidance in whole study and revised the manuscript. All authors read and approved the final manuscript.

## Acknowledgements

Thanks to the assistance of Mugu lnc (Beijing) in bioinformatics.

## Figure legends

**Figure S1**. Cell clustering. **(A)** Clustering of single-cell subpopulations. **(B)** The distribution of samples in clusters. **(C)** Annotation for all cell types.

**Figure S2. (A)** Heat map of the top five differential genes in each cluster. **(B)** B cell functional status analysis. **(C)** T cell functional status analysis. **(D)** Monocyte functional status analysis. **(E)** Macrophage functional status analysis. **(F)** NK cells functional status analysis. **(G)** Heat map of immune cell marker genes.

**Figure S3. (A)** Ligand-receptor interaction network diagram, different colours and sizes represent different numbers of node connections. **(B)** Multilayer network diagram—green is the ligand gene, red is the receptor gene, and yellow is the transcription factor.

**Figure S4**. WGCNA and functional analysis in black and magenta modules.

**(A)** Schematic diagram of threshold screening and determination. **(B)** Clustering dendrogram of all genes from last step. **(C)** Correlations of OS time with mean gene significance and errors in all modules. **(D)** Correlations of OS with mean gene significance and errors in all modules. **(E)** Heat map of the correlations between MEs and OS traits. **(F)** BP term in GO. **(G)** CC term in GO. **(H)** MF term in GO. **(I)** KEGG pathway analysis.

**Figure S5. (A-G)** Correlation analysis of risk score and other clinical factors. **(H-J)** The ROC curve was used to analyse the accuracy of risk scores and other clinical factors in predicting the 1-, 3-, and 5-year survival of patients with ccRCC.

**Figure S6. (A)** Cell infiltration analysis results. **(B)** The differential expression levels of genes in the risk prediction model obtained through the analysis of TCGA database. **(C)** The differential expression levels of genes in the risk prediction model obtained through the analysis of the CPTAC database. **(D)** Univariate association of the prognostic model and clinicopathological characteristics with overall survival. **(E)** Multivariate association of the prognostic model and clinicopathological characteristics with overall survival.

**Figure S7. (A)** Nomogram predicting 1-, 3- and 5-year OS for patients with ccRCC. **(B-D)** The calibration curve for predicting 1-, 3- and 5-year OS for patients with ccRCC. **(E-G)** Time-dependent ROC curve analysis evaluates the accuracy of the nomograms.

## Tables

**Table S1**. Notes on cell clustering.

**Table S2**. Differential genes in each cell cluster.

**Table S3**. Ligand-receptor relationship pair.

**Table S4**. Immune cell multifactor network relationship pair.

**Table S5**. Intersection genes in immune cell multifactor network relationship pair and TCGA.

**Table S6**. Genes in black and magenta models of WGCNA.

