## Supplementary figures and images for "Integrated analysis of single-cell and transcriptome based RNA-seq multilayer network and WGCNA for construction and validation of an immune cell-related prognostic model in clear cell renal cell carcinoma"

### Fig S1

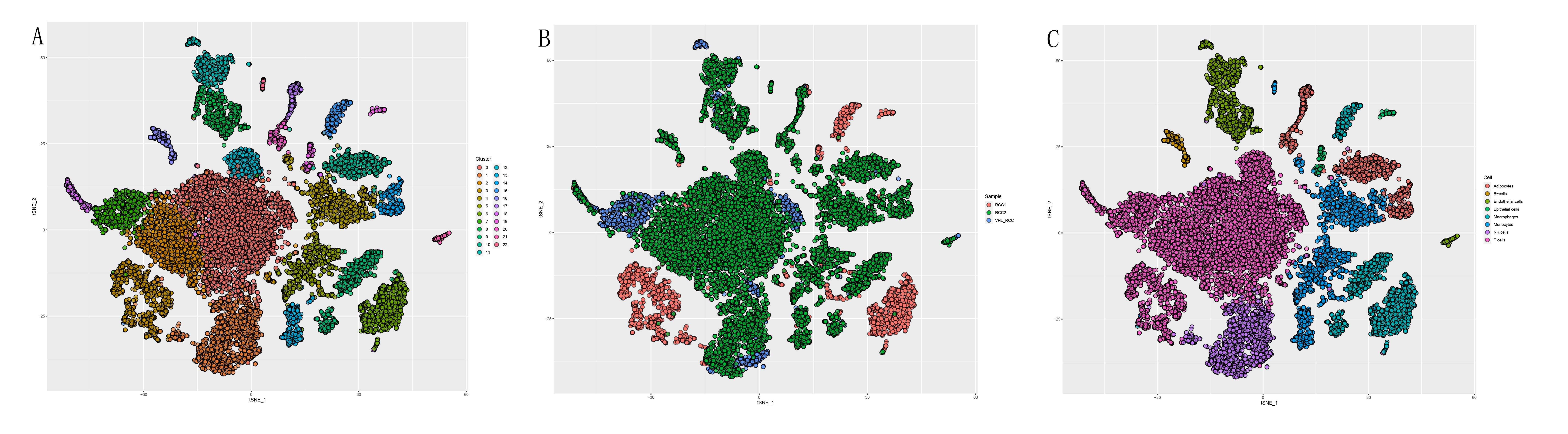

### Fig S2

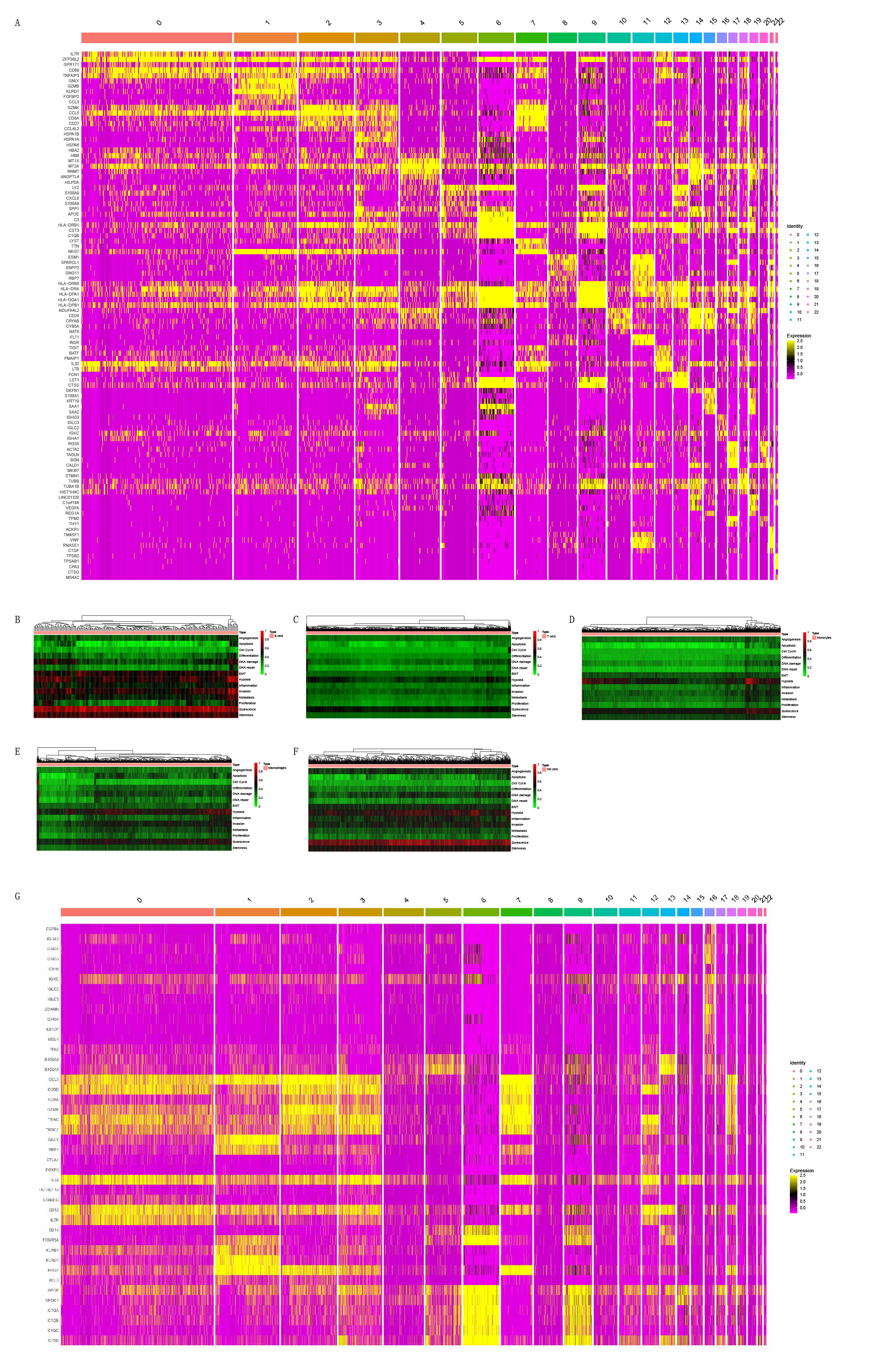

### Fig S3

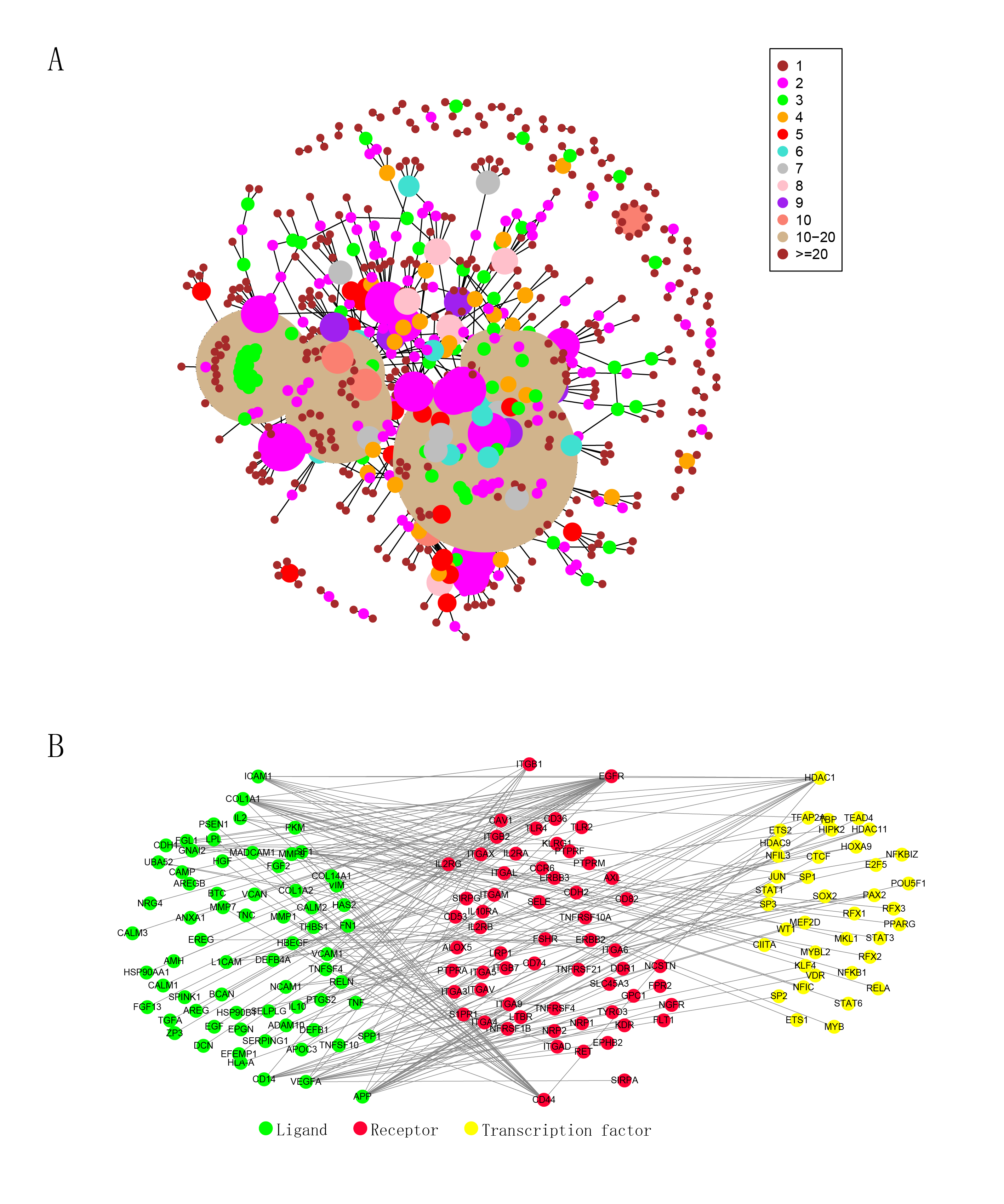

### Fig S4

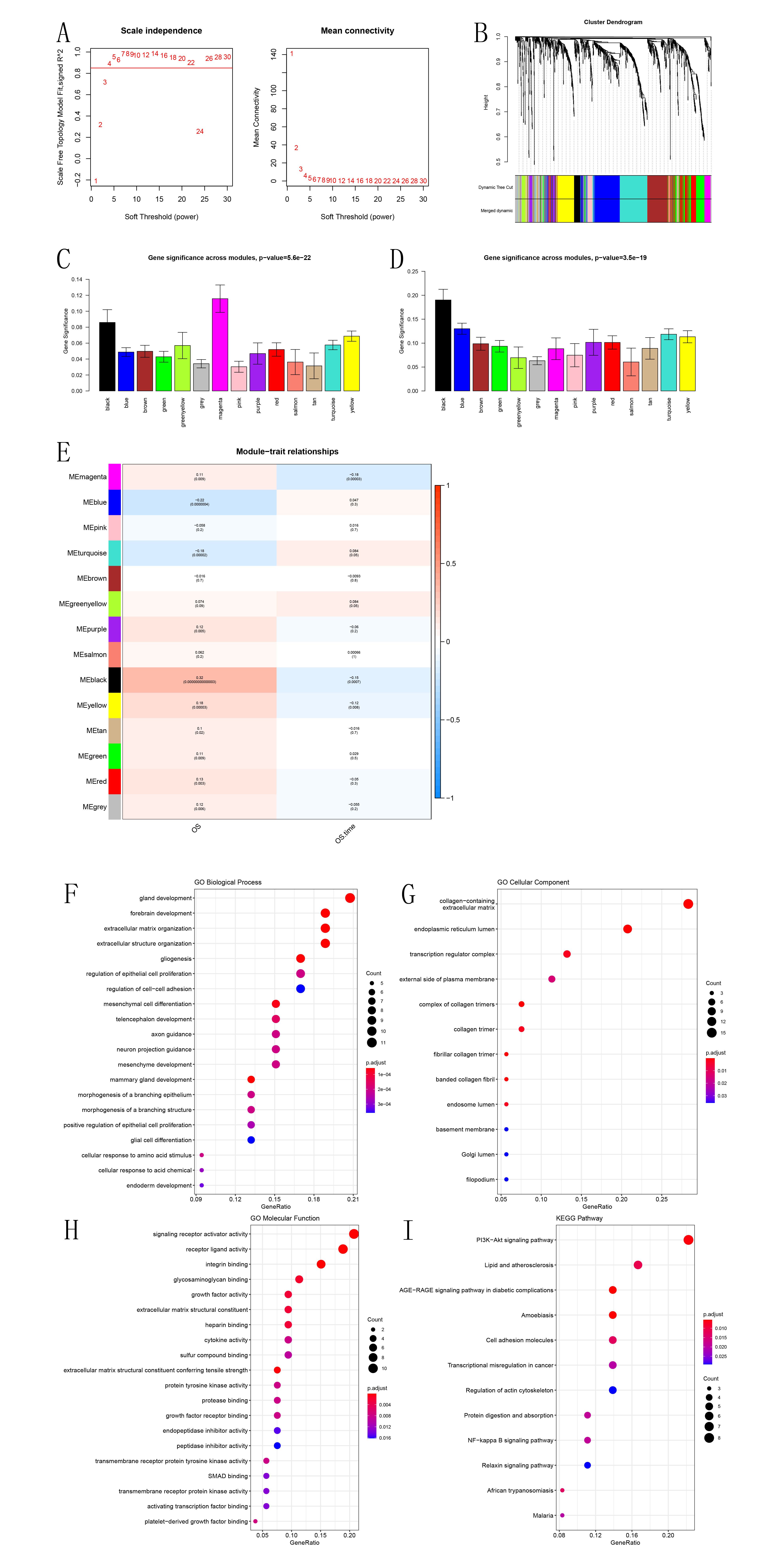

### Fig S5

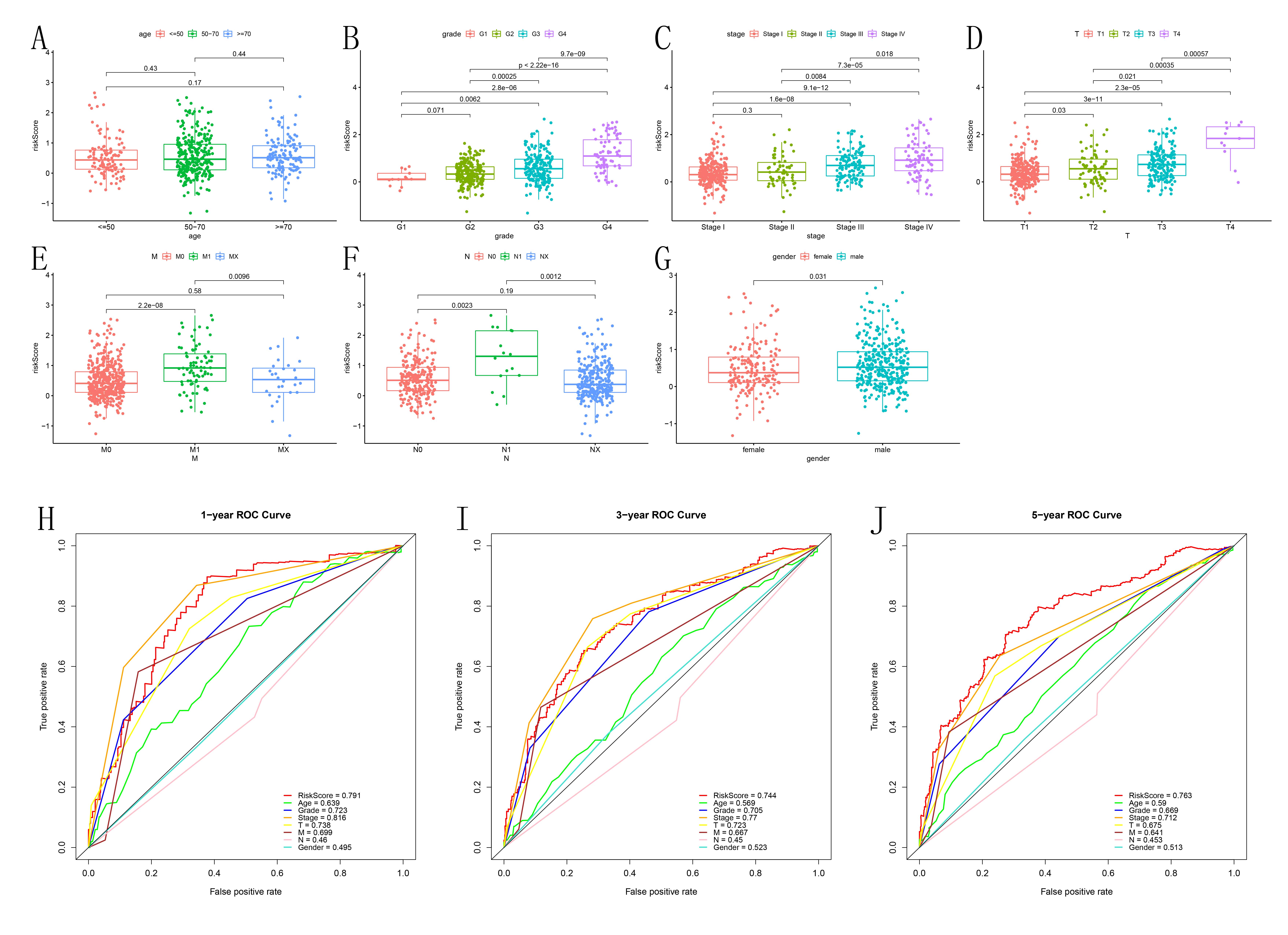

### Fig S6

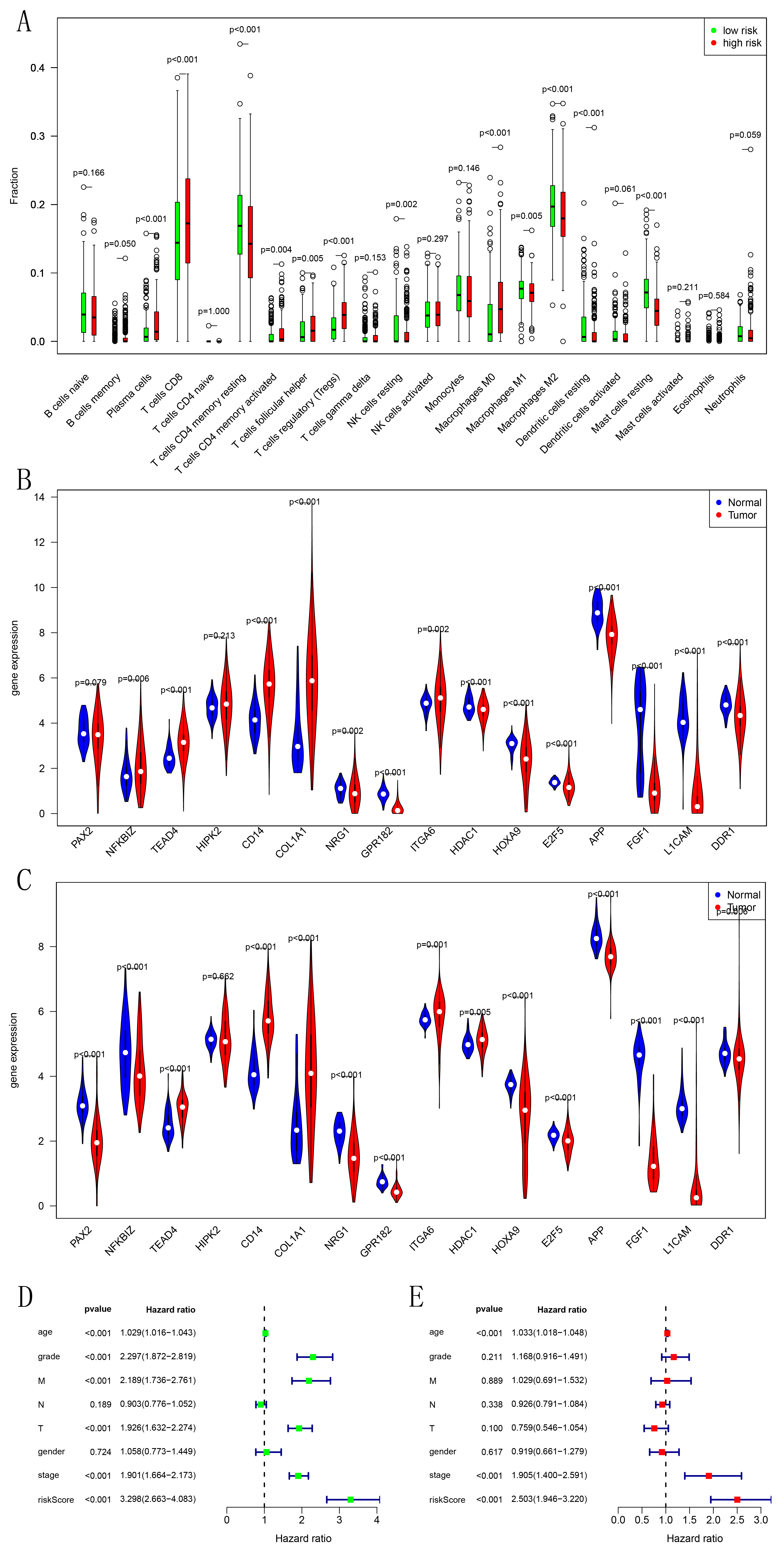

### Fig S7

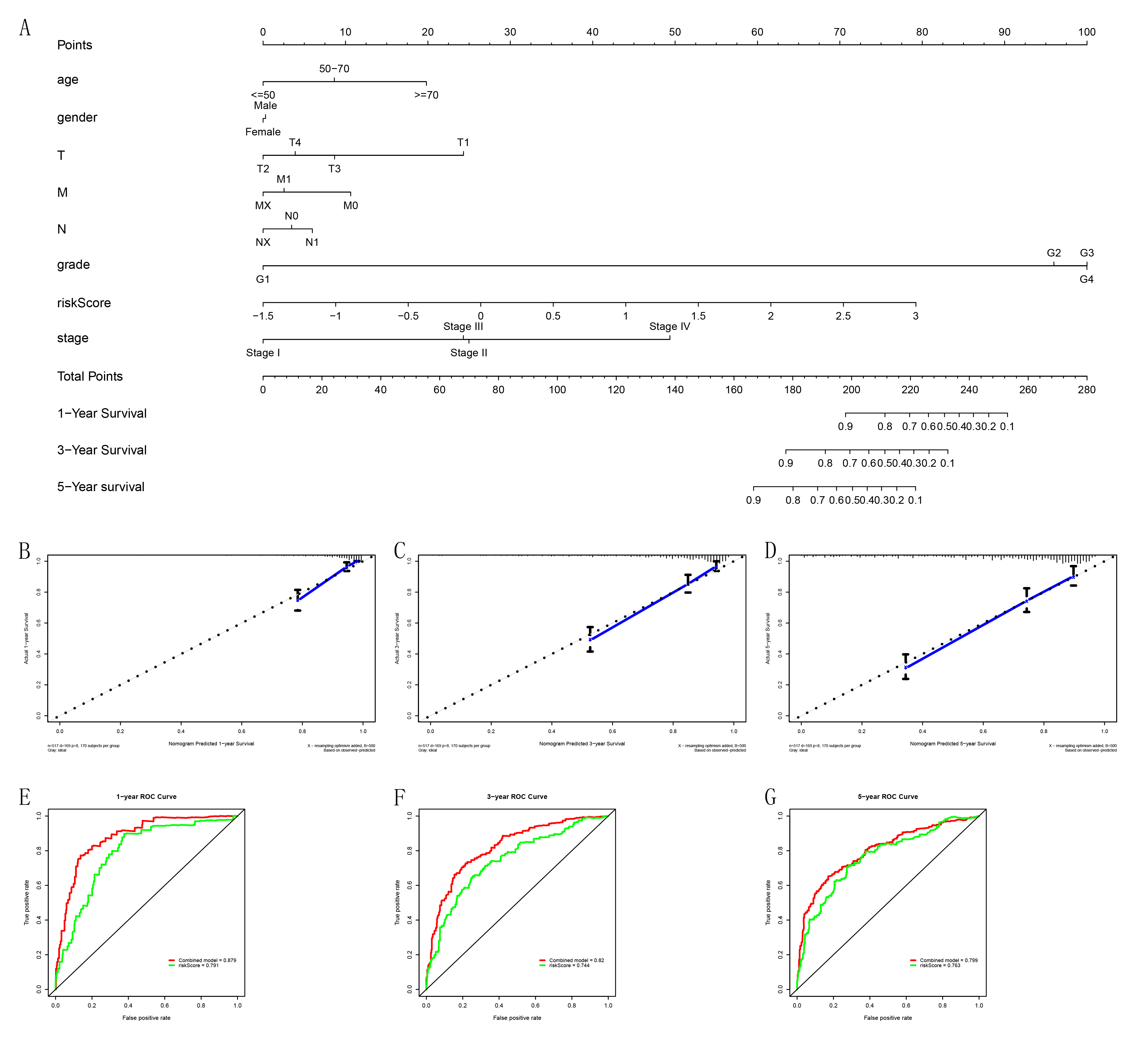
